# Genomic footprints of bottleneck in landlocked salmon population

**DOI:** 10.1101/2022.11.28.518181

**Authors:** Sankar Subramanian, Manoharan Kumar

## Abstract

At the end of the last ice age, several Atlantic salmon populations got caught up in the lakes and other small waterbodies of the Northern Hemisphere. Therefore, the pattern of evolution shaping the landlocked salmon populations is different from the other anadromous salmons, which migrate between the sea and rivers. According to the theories of population genetics, the effect of genetic drift is expected to be more pronounced in the former compared to the latter. Here we examined this using the whole genome data of landlocked and anadromous salmon populations of Norway. Our results showed a 50-80% reduction in the genomic heterozygosity in the landlocked compared to anadromous salmon populations. The number and total size of the runs of homozygosity (RoH) segments of landlocked salmons were 2 to 8-fold higher than those of their anadromous counterparts. We found the former had a higher ratio of nonsynonymous-to-synonymous diversities than the latter. The investigation also revealed a significant elevation of homozygous deleterious Single Nucleotide Variants (SNVs) in the landlocked salmon compared to the anadromous populations. All these results point to a significant reduction in the population size of the landlocked salmons, which might have started after the last glacial epoch. Previous studies on terrestrial vertebrates observed similar signatures of a bottleneck when the populations from Island and the mainland were compared. Since landlocked waterbody such as ponds and lakes are geographically analogous to Islands for fish populations, the findings of this study suggest the similarity in the patterns of evolution between the two.

## Introduction

The evolution of vertebrate populations in islands is quite different from that of those living on the mainland. Previous studies observed a reduction in the body sizes of island populations for most of the vertebrates, but an increase was reported in a few small animals (Benitez-Lopez, et al. 2021; Foster 1964). This was called Foster’s rule or the Island effect. While the dwarfism was owing to the limited resource availability, the absence of predators was thought to be the reason for gigantism. On the other hand, studies based on molecular markers such as allozymes, microsatellites, and mitochondrial sequence data found a reduction in heterozygosity in island populations compared to their mainland relatives (Boessenkool, et al. 2007; Dussex, et al. 2021; Frankham 1997; Hansson, et al. 2014; Jensen, et al. 2013; Palkopoulou, et al. 2015; Robinson, et al. 2016; White and Searle 2007). This could be due to the reduction in the effective population size that is forced by the limited availability of space in islands. The heterozygosity is determined by the effective population size and mutation rate (Crow and Kimura 1970). Since the mutation rate does not change significantly between the Island and mainland populations of the same species, the decline in the population size leads to a proportional reduction in the heterozygosity. Apart from heterozygosity, recent studies based on whole genome data found a much higher number of runs of homozygosity (RoH) segments in Island populations than their mainland counterparts (Dussex, et al. 2021; Palkopoulou, et al. 2015; Robinson, et al. 2019). For example, the RoH segments constitute 23% of the mammoth genome from Wrangel Island, but these segments comprise only 0.83% of the mainland European mammoth (Palkopoulou, et al. 2015).

The accumulation of deleterious mutations is also a hallmark of population bottleneck, and therefore a number of studies compared the deleterious mutation load between the Island and mainland populations (Dussex, et al. 2021; Johnson and Seger 2001; Palkopoulou, et al. 2015; Pedersen, et al. 2017; Robinson, et al. 2016; Robinson, et al. 2019; Woolfit and Bromham 2005). Most of these studies used the ratio of divergence or diversity at nonsynonymous and synonymous sites (dN/dS) to quantify the mutational load and found that this ratio was much higher in the island population than in their mainland counterparts. For instance, a previous study using 70 phylogenetically independent comparisons of populations from the island and mainland taxa showed that the dN/dS ratios of the former were significantly higher than those of the latter (Woolfit and Bromham 2005). Using whole genome data from human populations found a much higher proportion of deleterious SNVs in Greenland populations compared to mainland Europeans (Pedersen, et al. 2017). Furthermore, whole genome-based studies comparing Island and mainland fox (Robinson, et al. 2016) and kakapo (Dussex, et al. 2021) populations observed similar results. These studies also observed a much higher proportion of homozygous deleterious SNVs in Island populations than the mainland ones.

During the last ice age, salmon populations from the Northern Oceans were translocated to the water bodies such as ponds and lakes in Europe and Northern America and were eventually got locked up after the glacial epoch (Hutchings, et al. 2019; Kjaerner-Semb, et al. 2021; Tonteri, et al. 2005). The aquatic animal populations in landlocked waterbodies are isolated from their counterparts in the ocean. This scenario is similar to the isolation of terrestrial animal populations in Islands from their mainland populations. Hence the same Island rule will hold true for the landlocked salmon, and the forces of evolution shaping these salmon populations are expected to be similar. Therefore, we examined this by analysing the whole genome data from landlocked and anadromous salmon populations from Norway. We compared the whole genome heterozygosity, runs of homozygosity (RoH), and deleterious SNVs of landlocked and anadromous salmon populations.

## Results

### Genomic heterozygosity

The whole genome data from 29 salmons belonging to six populations of Norway were used to estimate the heterozygosity per site (see Supplementary Information). This revealed that the nucleotide diversity of the landlocked salmon population was 0.00037 (Figure 1). The diversity estimates for the anadromous salmons varied between 0.00056 to 0.00067. The genomic heterozygosity estimated for the landlocked salmon (Namsen_landlocked) population was significantly (at least, *P* < 0.0001, *Z* test) smaller than those observed for the five anadromous populations, including the one (Namsen_Bjoera) that was closely located to the former. The estimate for the anadromous Baltic Sea population was 50% higher than that of the landlocked population, and those obtained for other anadromous populations were 72% to 80% higher than that of the landlocked ones. Furthermore, except for the estimate of the Baltic Sea populations, the heterozygosity obtained for other anadromous populations was similar – statistically not different (*P* = 0.31).

**Figure 1.**
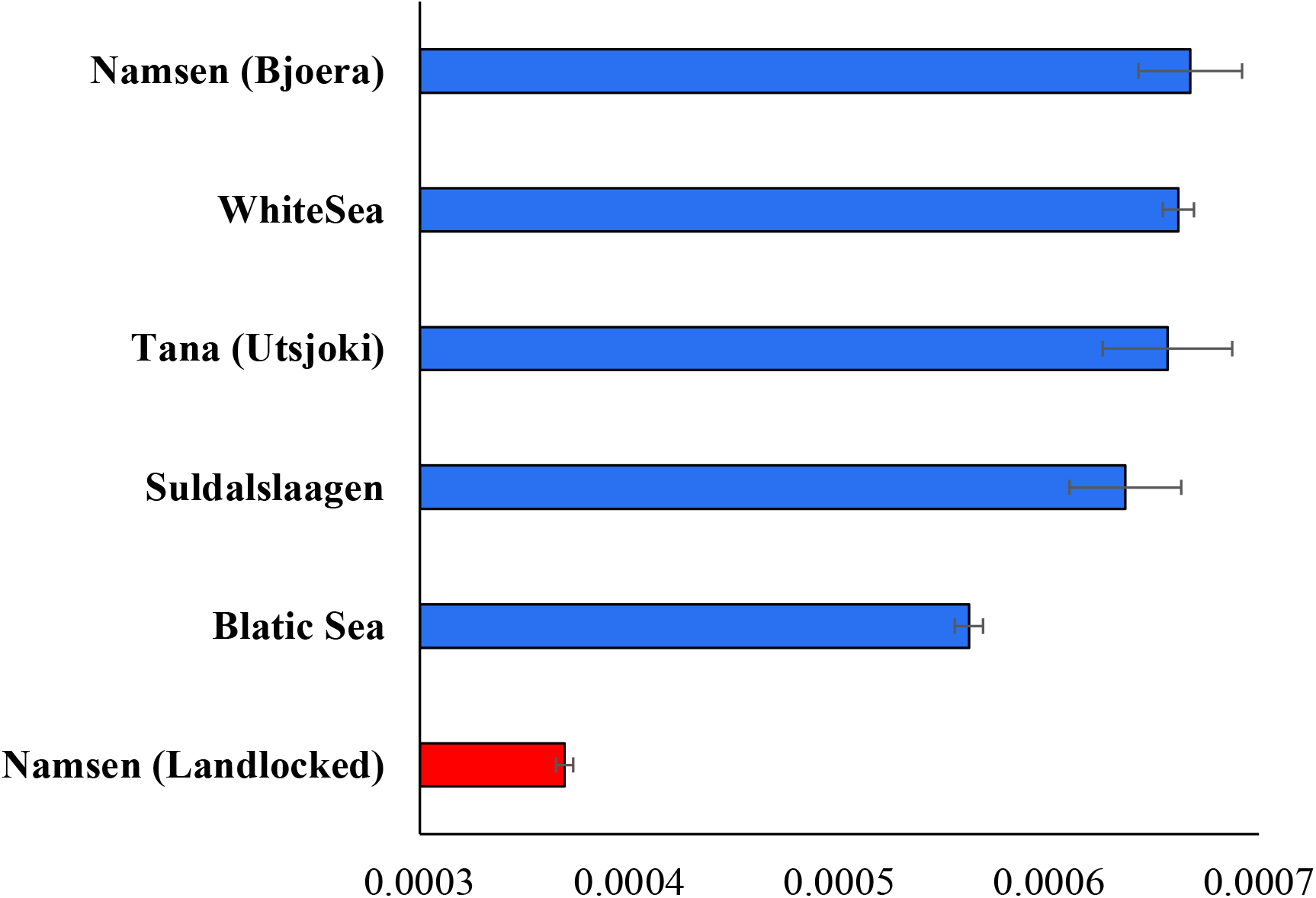
Genomic heterozygosity estimated for five anadromous and one landlocked population of Norway. Error bars denote the standard error of the mean. The genomic diversity of the landlocked salmon population was significantly smaller than those of anadromous populations (at least, *P* < 0.0001).

### Runs of homozygosity (RoH)

To understand the level of homozygosity between landlocked and anadromous salmon populations, we investigated the number and size of RoH in each genome. We used a threshold of >0.5 Mb to designate an RoH segment. Figure 2A shows that the mean number of RoH segments in landlocked salmons is 1180. The estimates for anadromous salmon populations range between 153 to 608. Therefore, the number of RoH segments in the landlocked population is approximately 1.9 to 7.7 times higher than those of the anadromous salmon populations, and the differences between them were highly significant (at least, *P* = 0.0003). We then compared the size of the RoH segments, which also revealed that the mean size of RoH segments in landlocked salmons was significantly (at least, *P* = 0.002) higher than those observed for anadromous salmons (Figure 2B). The estimate for the former was 393 Mb, and for the latter ranges between 46 to 205 Mb. Therefore, the mean size of RoH in landlocked salmons was approximately 1.9 to 8.5 times larger than those estimated for anadromous salmons.

**Figure 2.**
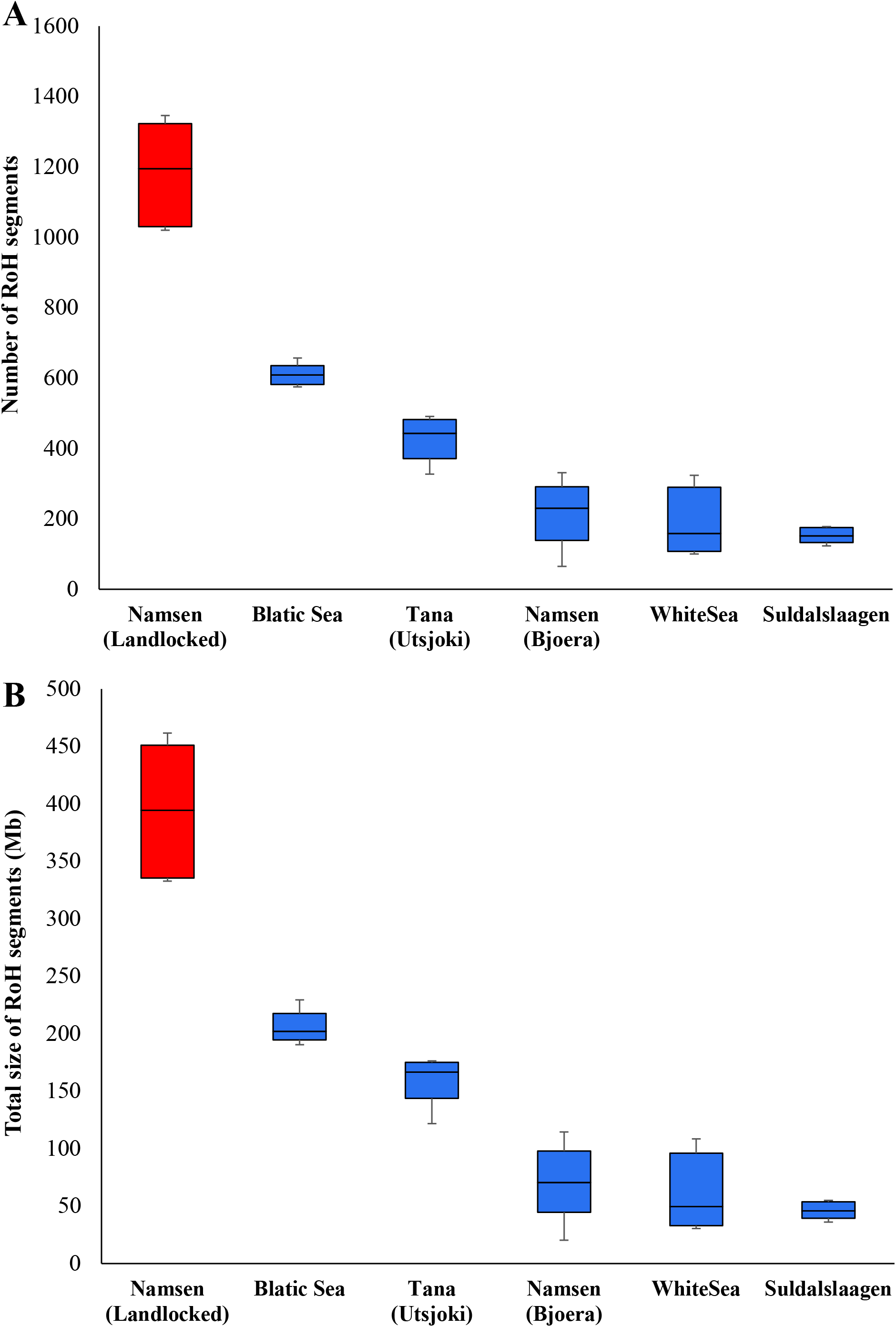
Box plot showing **(A)** the number and **(B)** the total size of runs of homozygous (RoH) segments in landlocked and anadromous salmon populations. The centre line denotes the median, the boundaries of the box represent the first and third quartiles and the whiskers show the maximum and minimum values. The mean number of RoH of the landlocked salmon population was significantly higher than those of anadromous populations (at least, *P* = 0.0003). Similarly, the average size of RoH of the landlocked salmon population was significantly higher than those of anadromous populations (at least, *P* = 0.002).

### The ratio of the diversities at nonsynonymous and synonymous sites

To measure the accumulation of deleterious variants, we first used the ratio of diversities at nonsynonymous and synonymous sites (dN/dS). For this purpose, we estimated the nucleotide diversities at amino acid changing and synonymous sites and computed their ratio. We then plotted the dN/dS ratio against the genomic heterozygosity for each individual (Figure 3A). The regression analysis revealed a highly significant (*r* = 0.79, *P* < 0.000001) negative correlation between the two variables. This suggests that individuals with low heterozygosities have high dN/dS ratios. Importantly, the dN/dS ratios of landlocked salmons appear to be high. This has been clarified in figure 3B, which shows that the mean dN/dS ratio estimated for landlocked salmons was significantly (P = 0.0018) higher than those observed for the anadromous salmon populations.

**Figure 3.**
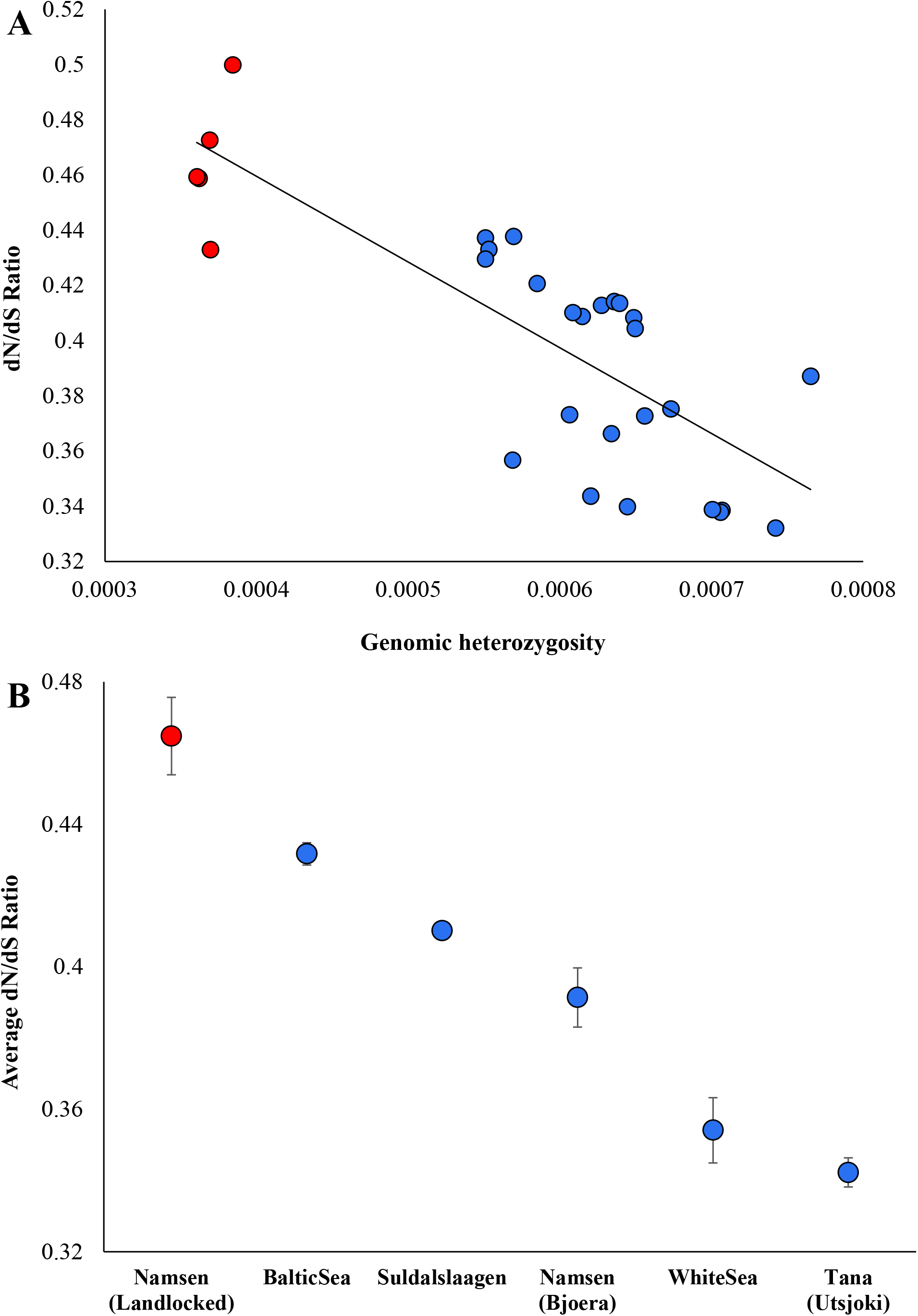
**(A)** The correlation between genomic diversity and the ratio of nonsynonymous and synonymous diversities (dN/dS). The relationship was highly significant (*r* = 0.79, *P* < 0.000001). The best-fitting regression line is shown. **(B)** The average dN/dS ratio was estimated for landlocked and anadromous salmon populations. Error bars show the standard error of the mean. The ratio observed for landlocked salmons was significantly higher than those of anadromous ones (at least, *P* = 0.0018).

We then calculated the number of homozygous and heterozygous nonsynonymous SNVs and plotted their proportions in stacked column graphs. Figure 4A reveals that the proportion of homozygous SNVs estimated for landlocked salmons was 60%, and for the anadromous salmons, this ranged between 42% to 48%. Furthermore, the average number of homozygous SNVs of the landlocked population (19,628) was significantly higher than those of anadromous populations (15,290 to 17,461) (at least, *P* < 0.0001). A similar analysis was performed using genome-wide deleterious SNVs, and we used the GERP score to identify deleterious SNVs (see methods). This showed a much higher proportion (68%) of homozygous deleterious SNVs in landlocked salmons compared to those estimated for anadromous populations (46% to 55%). The mean number of homozygous deleterious SNVs of the former (644) was significantly higher (at least, *P* < 0.0001) than those of the latter (457 to 536). Finally, the analysis using the loss of function (LoF) SNVs also revealed the same pattern. The landlocked populations had a much higher proportion of homozygous LoF SNVs (52%) than the proportions estimated for the anadromous populations (36% to 42%). As expected, the mean homozygous LoF SNV counts of the landlocked (1273) was significantly higher (at least, *P* < 0.0001) than those observed for the anadromous ones (954 to 1120).

**Figure 4.**
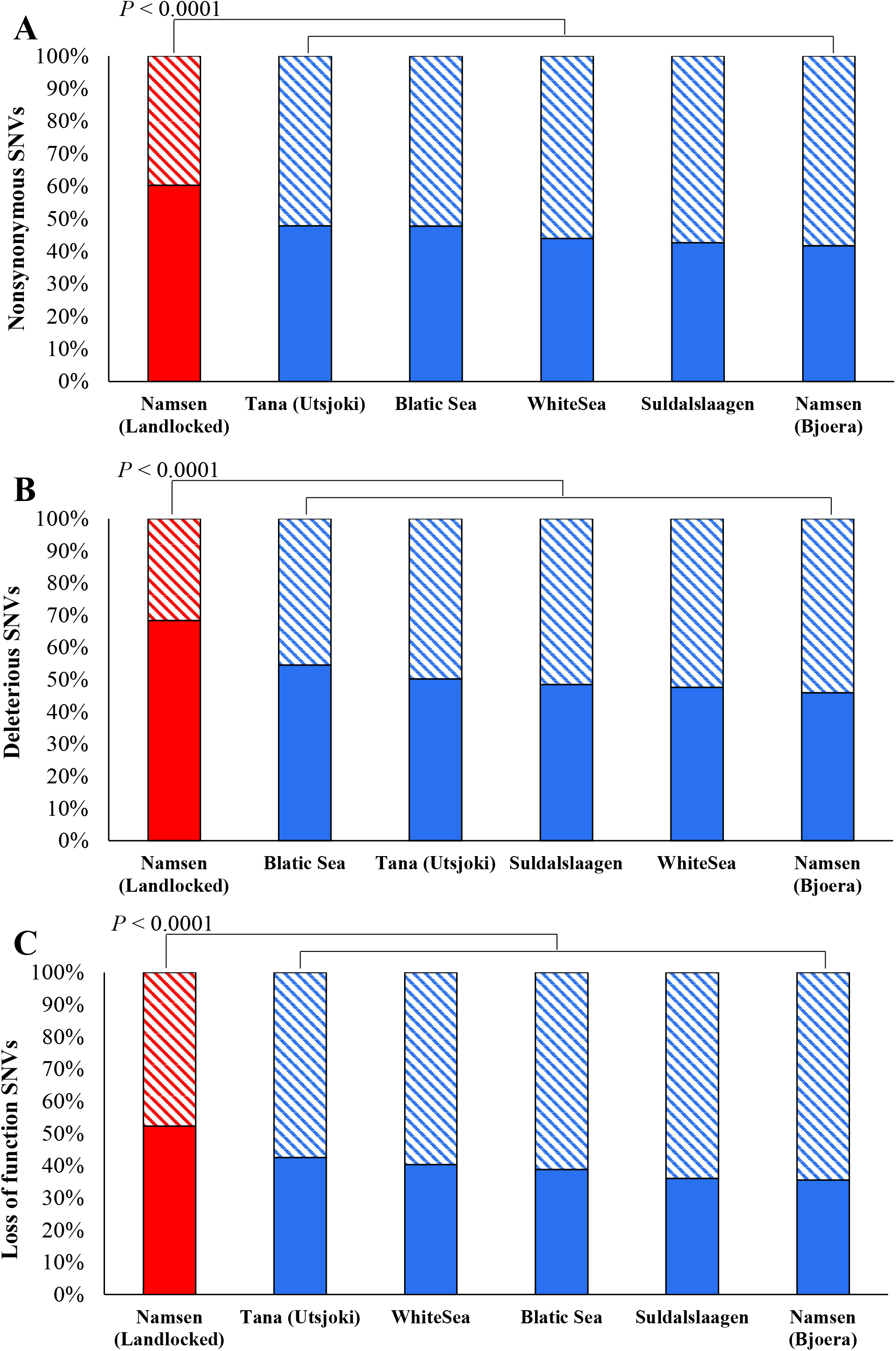
The stacked bar shows the proportions of homozygous (solid) and heterozygous (stripped) SNVs estimated for landlocked and anadromous salmon populations. **(A)** Nonsynonymous SNVs **(B)** Deleterious SNVs **(C)** Loss of function (LoF) SNVs. The number of homozygous SNVs in landlocked salmons was significantly higher than those estimated for the anadromous salmon, and this is true for the comparisons involving nonsynonymous, deleterious, and LoF SNVs (at least, P < 0.0001).

## Discussion

Using the whole genome data from anadromous and landlocked salmon populations, we performed comparative analyses, and our results provide four lines of evidence of severe bottleneck in the latter. First, we observed a much-reduced genomic diversity in landlocked populations compared to anadromous ones. The result from our whole genome data analysis confirms previous studies based on a few microsatellite and allozyme data from a few loci (Sandlund, et al. 2014; Vuorinen and Berg 1989). Furthermore, this result is similar to earlier studies on many terrestrial mammals and birds (Boessenkool, et al. 2007; Dussex, et al. 2021; Frankham 1997; Hansson, et al. 2014; Palkopoulou, et al. 2015; Pedersen, et al. 2017; Robinson, et al. 2016; Robinson, et al. 2019; White and Searle 2007). The heterozygosities estimated for the Island populations were 20% to 84-fold (e.g. Island fox) higher than those observed for their respective mainland counterparts (Robinson, et al. 2016). Second, the number and total size of RoH estimated for the landlocked genomes were 2 to 8-fold higher than those of anadromous populations. Earlier studies comparing Wrangel Island and European mainland mammoth populations showed a several-fold increase in the RoH content of the former (Palkopoulou, et al. 2015). Similar observation was reported comparing the Steward Island and mainland New Zealand populations of kakapo (Dussex, et al. 2021) and between the wolves of Isle Royale and mainland Minnesota (Robinson, et al. 2019). These studies showed that the Island populations had higher proportions of both medium-sized (0.2 to 1Mb) and long (>2Mb) RoH. While there was a significant number of medium-sized RoH in landlocked salmons (Figure 2), only a very few (<10) long RoH were observed. The latter suggests that there was no significant inbreeding in the landlocked populations. Because it has been shown that inbreeding produces long RoH, and on the contrary, the reduction in population size alone creates predominantly medium-sized RoH (Ceballos, et al. 2018).

Third, we showed a much higher dN/dS ratio for landlocked salmons than anadromous ones (Figure 3). The dN/dS ratio suggests the fraction of segregating (and fixed) nonsynonymous SNVs in comparison with the segregating neutral synonymous SNVs. Hence the higher ratio suggests proportionally higher nonsynonymous SNVs in the population. Since nonsynonymous SNVs could be deleterious, the dN/dS ratio was routinely used to measure the deleterious mutational load. A number of previous studies have shown a much higher dN/dS ratio in the Island populations of terrestrial vertebrates compared to their mainland relatives (Johnson and Seger 2001; Robinson, et al. 2016; Rogers and Slatkin 2017; Woolfit and Bromham 2005). Fourth, our findings revealed a much higher proportion of homozygous deleterious SNVs in landlocked salmons than in anadromous populations. This suggests that a greater number of heterozygous deleterious SNVs are converted to homozygous state due to strong genetic drift. In contrast, purifying selection prevents low-frequency SNVs to reach high frequencies (Kimura 1983) and hence heterozygous SNVs are not allowed to be converted to homozygous ones. Similar patterns were reported in Island foxes (Robinson, et al. 2016), Isle Royale wolves (Robinson, et al. 2019), and Greenland Inuit populations (Pedersen, et al. 2017) in comparison with their respective mainland cousins.

All results of this study point out that there was a significant reduction in the population size of the landlocked salmons after they had been captured in the land-encircled section of the Namsen river with a surface area of 12 km^2^. This is similar to those observed for the waterlocked Island populations of terrestrial vertebrates. Therefore, we can predict that the pattern of evolutionary forces shaping the landlocked populations will be very similar to those operating on the terrestrial vertebrate populations living on the Islands.

## Materials and Methods

### Genome data

The whole genome raw sequence data in the fastq format for five landlocked salmons were available from a previous study (Bertolotti, et al. 2020). For comparison, we obtained the whole genome data for 24 anadromous salmons belonging to five locations that were selected in order to include those representing a wide geographic area of Norway. The landlocked salmons were from an enclosed part of the river Namsen with a surface area of 12 km^2^. For comparison, we included five anadromous salmons from the Namsen river that was open to the sea. Additionally, five anadromous salmons from Southern Norway (Suldalslaagen river), five from Northern Norway (Tana river), five from the Baltic Sea, and four from the White Sea were included. Furthermore, we included the reference genome data (Sal_tru 1.1) of river trout (*Salmo trutta*) to use as the outgroup (see Supplementary Information).

### Bioinformatic data processing

The *fastq* reads from 29 complete genomes were mapped to the Salmon reference genome (build Ssal v3.1) using the *bwa* aligner (Li and Durbin 2009). The mapped reads in the sequencing alignment mapped (SAM) format was converted to binary alignment/mapped (BAM) format using *Samtools* (Li, et al. 2009). The aligned reads were then sorted based on the chromosomal positions, and then the PCR duplicates were removed using the *Picard* tool (https://broadinstitute.github.io/picard/). The genotypes for all chromosomal positions were called using *Samtools*. All 29 vcf files were merged into one file, and this was done for each chromosome. Finally, we filtered biallelic variant sites using an in-house awk script, which resulted in 25 million SNVs. In addition, we also estimated the average read depth for each sample using the “depth” module of the software *Samtool*. The total number of sites covered was calculated for each sample using an in-house script. The number of runs of homozygosity segments was estimated using the *plink* software (Purcell, et al. 2007) with following parameter (--geno 0.01 --homozyg --homozyg-window-het 0 --maf 0.05). We used the whole genome data of the river trout to determine the direction of mutational change and to identify the derived alleles.

### Population genetic analysis

To annotate protein-coding regions and to identify the functional consequences of SNVs, the SNVs tool *SNPeffect* was used (De Baets, et al. 2012). Based on the annotations, we identified synonymous and nonsynonymous SNVs. To determine deleterious SNVs, we used the GERP scores (Cooper, et al. 2005), which were obtained from the data resource server Ensembl. The GERP score for each chromosomal position was calculated using a multiple sequence alignment containing 90 fish genomes (https://ftp.ensembl.org/pub/release-108/bed/ensembl-compara/65_fish.gerp_constrained_element/gerp_constrained_elements.salmo_salar.bb). We used a threshold of >4 to designate an SNV to be deleterious in nature. Using the genome annotations, we also identified the highly deleterious SNVs that cause premature termination or loss of function (LoF) of proteins. The keywords “stop_lost”, “stop_gained”, “start_lost”, “splice_donor” and “splice_acceptor” were used to detect the LoF SNVs.

The numbers of synonymous and nonsynonymous SNVs were divided by their respective number of synonymous and nonsynonymous sites to obtain the diversities at these sites. The ratio of the two was used to determine the accumulation of deleterious nonsynonymous SNVs in each genome. The average number of LoF and deleterious SNVs per genome, along with the standard errors, were estimated for each genome. The significance between the mean counts was determined using the Z test, and the statistical significance was determined using the software Z to P (http://vassarstats.net/tabs_z.html). A Pearson correlation coefficient was used to determine the strength of the correlation. The statistical significance of the correlation was determined by converting the correlation coefficient *r* to the normal deviation Z, and this was accomplished using the online software *r* to *P* (http://vassarstats.net/tabs_r.html).

## Supplementary Material

Supplementary data are available at Genome Biology and Evolution online.

## Data Availability

Raw read sequence data used in this study was obtained from the SRA database. The details of the accession numbers and metadata are given in the supplementary material.

